# TRPV4 is the temperature-sensitive ion channel of human sperm

**DOI:** 10.1101/268409

**Authors:** Nadine Mundt, Marc Spehr, Polina V. Lishko

**Affiliations:** Department of Molecular and Cell Biology, University of California, Berkeley, CA 94720, USA; Department of Chemosensation, Institute for Biology II, RWTH Aachen University, D-52074 Aachen, Germany

**Author notes:** corresponding author Polina Lishko < >.

**Keywords:** TRPV4, sperm ion channels, hyperactivation, cation channel, thermosensitivity

## Abstract

Ion channels control sperm fertilizing ability by triggering hyperactivated motility, which is regulated by membrane potential, intracellular pH, and cytosolic calcium. Previous studies unraveled three essential ion channels that regulate these parameters: 1) the Ca^2+^ channel CatSper, 2) the K^+^ channel KSper, and 3) the H^+^ channel Hv1. However, the molecular identity of an additional sperm Na^+^ conductance that mediates initial membrane depolarization and, thus, triggers downstream signaling events is yet to be defined. Here, we functionally characterize DSper, the Depolarizing Channel of Sperm, as the temperature-activated channel TRPV4. It is functionally expressed at both mRNA and protein levels, while other temperature-sensitive TRPV channels are not functional in human sperm. DSper currents are activated by warm temperatures and mediate cation conductance, that shares a pharmacological profile reminiscent of TRPV4. Together, these results suggest that TRPV4 activation triggers initial membrane depolarization, facilitating both CatSper and Hv1 gating and, consequently, sperm hyperactivation.

## Introduction

The ability of human spermatozoa to navigate the female reproductive tract and eventually locate and fertilize the egg is essential for reproduction [1]. To accomplish these goals, a spermatozoon must sense the environment and adapt its motility, which is controlled by ATP production and flagellar ion homeostasis [2]. Of vital importance is the transition from symmetrical basal tail bending into “hyperactivated motility” – an asymmetrical, high-amplitude, whip-like beating pattern of the flagellum - that enables sperm to overcome the egg’s protective vestments. The steroid hormone progesterone (P4) acts as a major trigger of hyperactivation[3], [4]. P4 is released by cumulus cells surrounding the egg [5] and causes a robust elevation of sperm cytoplasmic [Ca^2+^] via the principal Ca^2+^ channel of sperm, CatSper (EC_50_= 7.7 ± 1.8 nM) [6]–[8]. The steroid acts via its non-genomic receptor ABHD2, a serine hydrolase that, upon P4 binding, releases inhibition of CatSper [9]. The propagation of a Ca^2+^ wave produced from the opening of CatSper channels along the flagellum is a necessary milestone in the process of fertilization and initiates hyperactivated motility [10]. CatSper channels exhibit pronounced voltage-dependency (slope factor k ~ 20 in humans) with half-maximal activation at V_1/2 human_ = +70 mV [6]. Given this unusually high V_1/2_, only a small fraction of human CatSper channels are open at physiological negative resting membrane potentials. P4 has been shown to potentiate CatSper activity by shifting V_1/2_ to more negative values (V_1/2 human_ = +30 mV with 500 nM P4 [6]). However, CatSper still requires both additional intracellular alkalization and significant membrane depolarization to function properly [2], [11]. The proton channel Hv1 was revealed as one of the regulators of intracellular pH (pH_i_) in human spermatozoa [11], [12]. By mediating unidirectional flow of protons to the extracellular environment, voltage-gated Hv1 represents a key component in the CatSper activation cascade, but it also induces membrane hyperpolarization by exporting positive charges out of the cell. Hv1 is also voltage-operated and requires membrane depolarization to be activated [11] Therefore, both CatSper and Hv1 must rely on yet unidentified depolarizing ion channels. P4 was shown to inhibit the K^+^ channel of human sperm KSper (IC_50_=7.5 ± 1.3 μM [13], [14]) making KSper one of the potential origins for membrane depolarization. However, efficient KSper inhibition requires P4 concentrations in the μM range, which are only present in close vicinity of the egg. Sperm hyperactivation, however, occurs in the oviduct, where P4 concentrations are not sufficient to block KSper [15]. Hence, the current model is missing a fourth member – the “Depolarizing Channel of Sperm” (DSper) [16]. Activation of this hypothetical DSper would induce long-lasting membrane depolarization and provide the necessary positive net charge influx for CatSper/Hv1 activity. Despite its central role, the molecular identity of DSper yet remains elusive.

The goal of this work was to characterize DSper and resolve its molecular identity in human spermatozoa. Using whole-cell voltage-clamp measurements, we recorded a novel non-CatSper conductance in both capacitated and noncapacitated spermatozoa. This unidentified, nonselective cation conductance exhibited outward rectification and pronounced temperature sensitivity. Parallel patch-clamp and Ca^2+^ imaging recordings demonstrated that the specific TRPV4 channel agonist RN1747 potentiated both DSper currents and intracellular calcium levels. Based on electrophysiological, biochemical and immunocytochemical data, we thus conclude that the molecular correlate of DSper is TRPV4.

## Results

### A novel non-CatSper conductance

As many calcium channels, CatSper conducts monovalent ions, such as Cs^+^ and Na^+^ in the absence of divalent cations from the extracellular solution (divalent free; DVF) [17]. CatSper is also permeable to Ca^2+^ and Ba^2+^, but it cannot conduct Mg^2+^ (Fig. 1 – Suppl. Fig. 1). In presence of extracellular Mg^2+^ the CatSper pore is blocked, resulting in the inhibition of monovalent CatSper currents (*I*_CatSper_) (Suppl. Fig. 1).

In whole-cell voltage-clamp recordings from human ejaculated spermatozoa, we consistently observed residual currents when *I*_CatSper_ was blocked with 1 mM extracellular Mg^2+^ (Fig. 1 A, B). Cs^+^ inward currents elicited under DVF condition (black traces and bars) were larger than currents recorded in the presence of Mg^2+^ (red traces and bars) (Fig. 1 A-C). This phenomenon was observed in both noncapacitated and capacitated spermatozoa, respectively. Notably, capacitated cells generally showed increased current densities under both conditions (Fig. 1 C). The data suggested that the remaining conductance is a novel non-CatSper conductance via the yet to be identified DSper ion channel. DSper currents were potentiated during capacitation (Fig. 1 C) and exhibited outward rectification. Though, DSper current recorded from capacitated cells was notably less rectificatying (Fig. 1 A, B). This DSper component is unlikely a remnant of an increased leak current, since the cells returned to their initial “baseline” current after returning to initial (HS) bath solution (Fig. 1 - Suppl. Fig. 2). Cation influx is the physiologically relevant entity to be analyzed, because it represents channel activity under physiological relevant conditions and ensures membrane depolarization. Therefore, from now on we analyzed DSper inward currents elicited by the change of membrane potential from 0 mV to −80 mV. To rule out ‘contamination’ of putative *I*_DSper_ with remaining *I*_CatSper_, we next tested whether 1 mM Mg^2+^ is sufficient to completely block *I*_CatSper_ and selectively isolate DSper currents. The CatSper inhibitor NNC 55-0396 [6] did not elicit any significant inhibitory effect on *I_DSper_* (Fig. 1 D-F), confirming efficient CatSper pore block by Mg^2+^. These findings corroborate our hypothesis that a novel CatSper-independent cation conductance could provide additional depolarization under physiological conditions. To isolate *I*_DSper_, we performed all following experiments in presence of both Mg^2+^ and NNC 55-0396.

**Figure 1:**
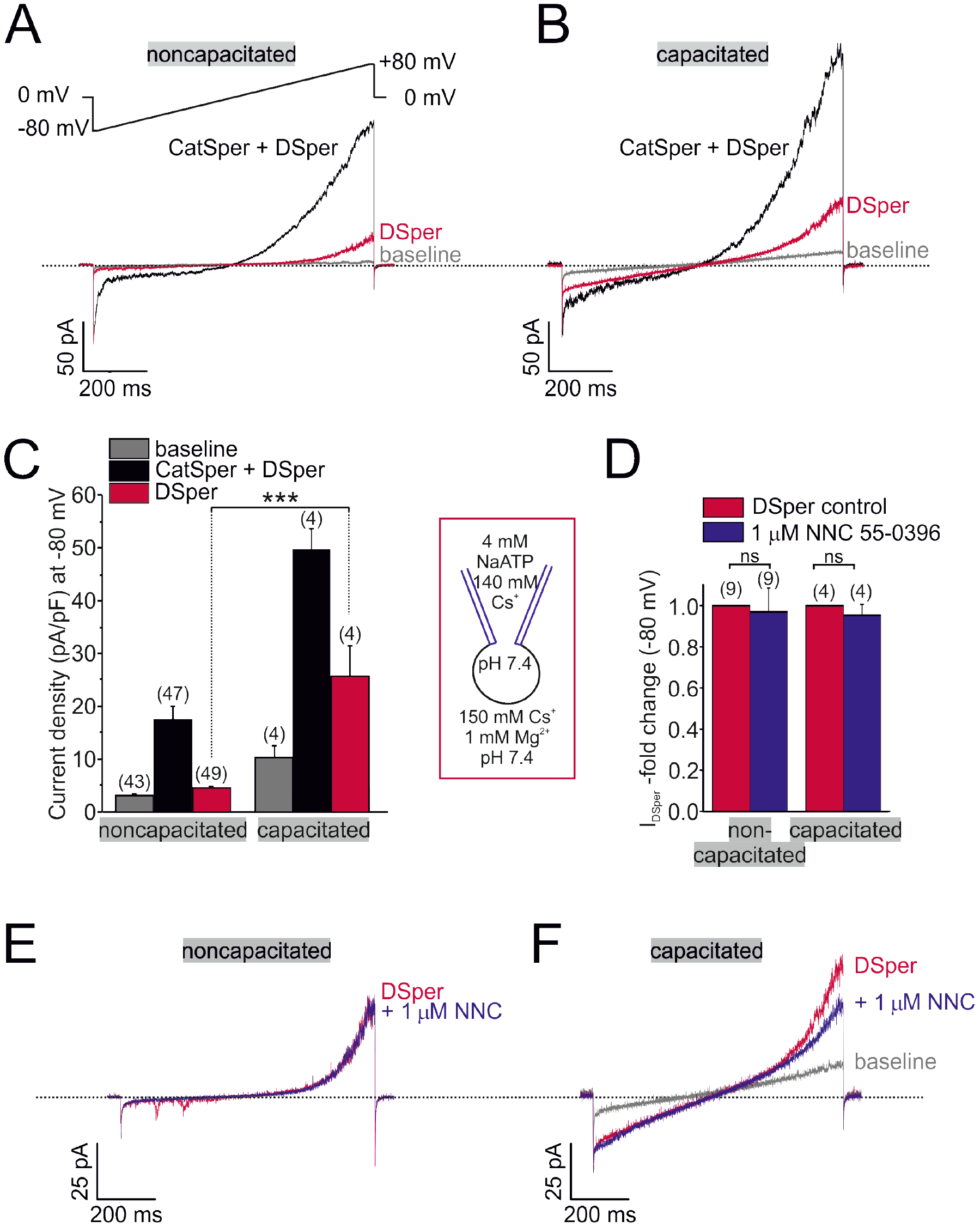
Electrophysiological recordings reveal a novel non-CatSper conductance. (A-B) Original current traces from representative whole-cell patch-clamp recordings from noncapacitated (A) and capacitated (B) human spermatozoa. Inward- and outward currents were elicited with voltage ramps as depicted in (A). Under divalent free conditions (black traces), typical CatSper monovalent caesium currents can be recorded. In presence of 1 mM Mg^2+^ (red traces), an outward rectifying “DSper” current component remains. Hence, the black traces represent a mixture of both CatSper and DSper monovalent Cs^+^ currents, while the red traces show pure Cs^+^ currents through DSper. (C) Quantification of current densities for all three conditions in A-B. DSper inward currents are potentiated upon capacitation (noncapacitated cells: −4.50 ± 0.41 pA/pF (n = 49), capacitated cells: −25.58 ± 5.88 pA/pF (n = 4). Statistical significance (unpaired t-test) was indicated by: \*\*\**p ≤0.001*. No variation between human donors were noticed. Quantification of normalized DSper inward currents (D) and original current traces (E-F) in presence and absence of the CatSper inhibitor NNC 55-0396 demonstrate the absence of inhibition.

### DSper current exhibits temperature sensitivity

We next aimed to investigate mechanism(s) of DSper activation mechanism. Previous work had focused on various DSper candidates, one being ATP-activated P2X channels. Navarro *et al.* showed functional expression of P2×2 in mouse spermatozoa [18]. However, human spermatozoa appear to be insensitive to extracellular ATP [19]. De Toni *et al*. suggested that human spermatozoa perform thermotaxis mediated by a member of the thermosensitive transient receptor potential vanilloid channel family, TRPV1 [20] supporting their claim by immunocytochemistry and Ca^2+^ imaging. By contrast, Kumar *et al.* detected TRPV4 expression in human spermatozoa using immunocytochemistry [21], yet the channel localization was not flagellar. Since the functional expression of a thermosensitive TRPV isoform in human spermatozoa is currently under debate, and their cation permeability renders them DSper candidates, we investigated the impact of temperature on DSper activity. As shown in Figure 2 A-C, elevating temperature profoundly increased *I*_DSper_. We observed a temperature-induced potentiation of both inward and outward currents in noncapacitated, as well as capacitated spermatozoa. A temperature ramp from 23 °C to 37 °C potentiated *I*_DSper_ inward currents by factors of 2.7 ± 0.5 for noncapacitated cells and 2.0 ± 0.2 for capacitated cells, respectively (Fig. 2 D). Half-maximal activation was achieved at T_1/2_ = 33 °C (noncapacitated) and T_1/2_ = 32 °C (capacitated) (Fig. 2 D). Moreover, the temperature-induced potentiation effect was reversible for both noncapacitated and capacitated cells (Figure 2 E). We hence conclude that the observed phenomenon is not a temperature-induced loss of the seal and compromised membrane stability and that DSper is indeed temperature-activated.

**Figure 2:**
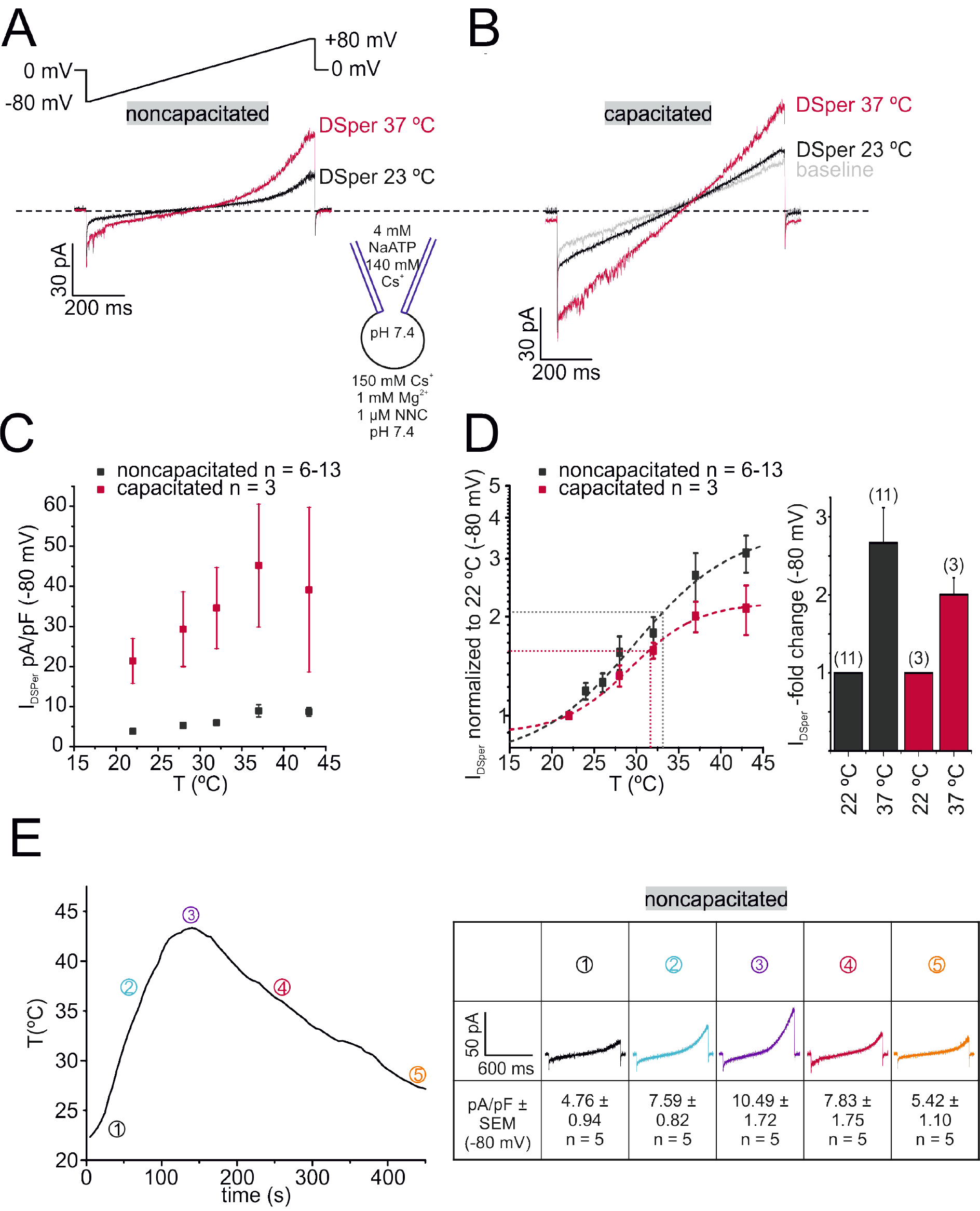
DSper is activated by warm temperatures. (A-B) Representative current traces from whole-cell patch-clamp recordings from noncapacitated (A) and capacitated (B) human spermatozoa challenged with a rise in temperature from 24 °C to 39 °C. Both DSper inward- and outward currents are increased at higher temperature. (C) Quantification of DSper inward current densities as a function of bath temperature (in °C). Noncapacitated (black squares) as well as capacitated cells (red squares) show increased current densities when stimulated with increasing bath temperatures. (D) Data of (C) normalized to room temperature (22 °C). Half maximal activation at T_1/2_ = 33 °C (noncapacitated) and T_1/2_ = 32 °C (capacitated) indicated by the dotted lines. (E) Mean applied bath temperatures as a function of time and corresponding DSper currents. Inset shows representative traces indicating that the temperature-induced potentiation effect was reversible.

### DSper conducts Na^+^

Since Na^+^ is the major permeant extracellular ion in the female reproductive tract ([Na^+^] = 140 − 150 mM [22]), Na^+^ is a likely source for membrane depolarization. We therefore investigated whether DSper has the capacity to conduct Na^+^. As indicated in Figure 3A, a similar outward rectifying *I*_DSper_ was recorded when extracellular Cs^+^ was replaced with equimolar concentrations of Na^+^. *I*_DSper_ inward Na^+^ currents were entirely CatSper-independent, since NNC 55-0396 had no significant inhibitory effect (Fig. 3B, C). In presence of both 1 mM Mg^2+^ and 1 μM NNC 55-0396, *I*_DSper_ was still reversibly activated by warm temperatures (Fig. 3 D, E) with a 4.1 ± 0.5 -fold increase for inward currents from 220C to 37CC. Half -maximum activation was at T_1/2_ = 32°C, comparable to previously analyzed values for temperature-activated Cs^+^ currents. Together, these electrophysiological data indicate that DSper shares characteristic hallmarks with thermosensitive TRPV channels [23]. We thus proceeded to define which TRPV channel(s) is involved.

**Figure 3:**
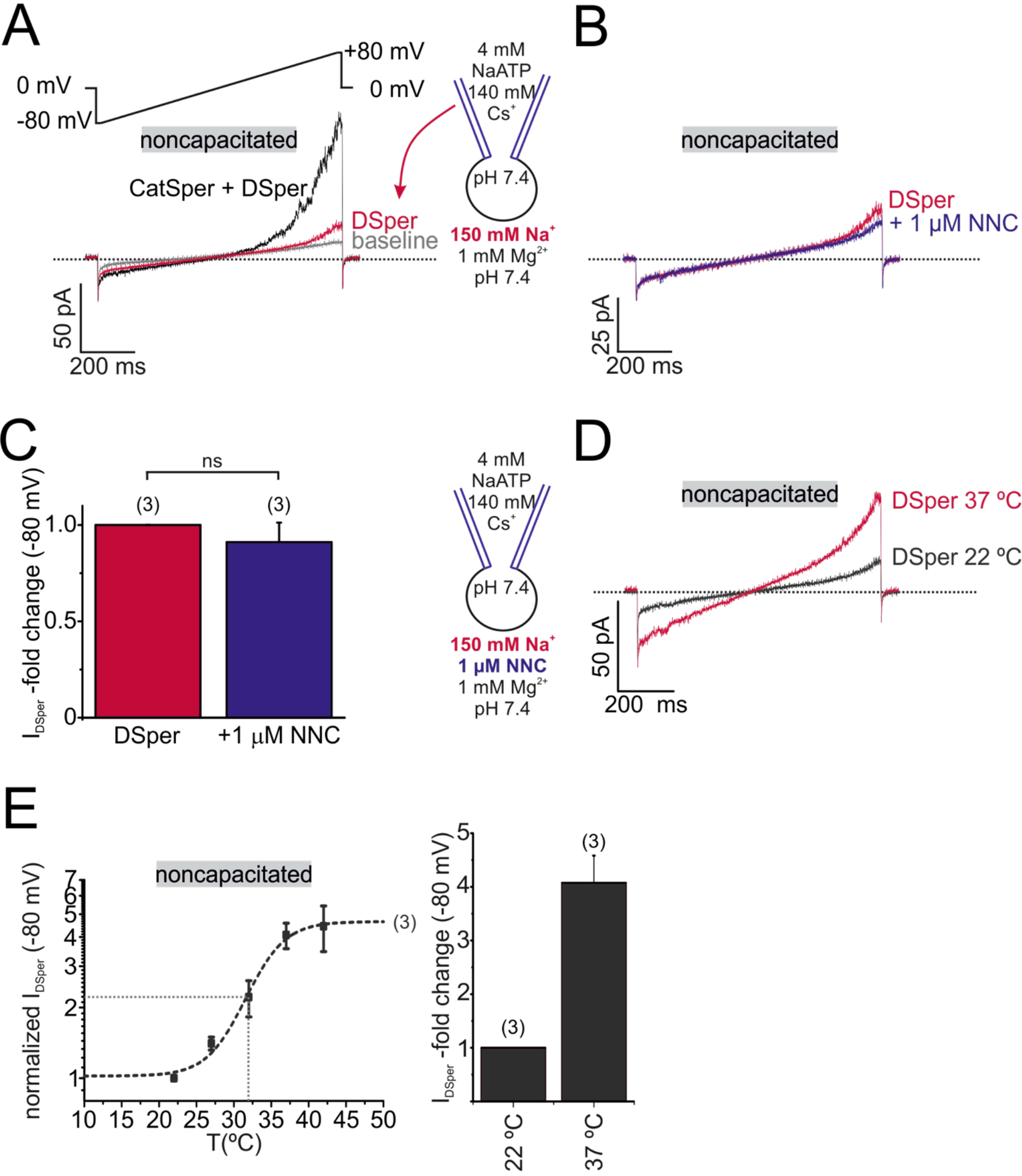
DSper conducts sodium. (A) Representative current traces from whole-cell patch-clamp recordings of noncapacitated human spermatozoa. Inward- and outward currents were elicited with voltage ramps as depicted. To record DSper currents, extracellular Cs^+^ was substituted with the same concentration of sodium Na^+^. Representative current traces (B) and quantification of normalized DSper inward currents (C) before and after stimulation with 1 μM NNC suggest that CatSper does not contribute to the recorded sodium inward conductance. (D-E) Representative current traces in (D) and normalized inward currents (E) at increasing bath temperatures. A similar temperature-induced potentiation effect of DSper sodium inward currents can be described as for caesium currents. Half maximal activation was achieved at T_1/2 sodium_ = 32 °C (dotted line).

### DSper is represented by the cation channel TRPV4

Based on the observed *I*_DSper_ temperature spectrum (Fig. 2D and 3E), candidate channels are TRPV3 and TRPV4 [24]. In addition, TRPV1 was previously proposed as a mediator of human sperm thermotaxis [20]. To discriminate between these channels, we tested potential effects of corresponding selective agonists – carvacrol [25], RN1747 [26] and capsaicin [27]. Employing both electrophysiological and Ca^2+^ imaging recordings, only the TRPV4 agonist RN1747 showed an effect. In detail, application of 1 μM RN1747 (EC_50_ = 0.77 μM [26]) potentiated DSper inward and outward currents (Fig. 4 A, B). Potentiation of DSper outward currents was more prominent than for DSper inward currents (factor 1.42 ± 0.17 for inward currents, factor 1.95 ± 0.35 for outward currents; Fig. 4 C). Yet, one can easily see an increase of the inward current noise factor that may indicate single-channel opening (Fig. 4B). Using Ca^2+^ imaging in fluo-4/AM-loaded sperm (Video 1, Video 2, Fig. 4), we next recorded fluorescence changes in the flagellar principle piece (Fig. 4D, top) while stimulating with 1 μM RN1747. Applicaton of the TRPV4 agonist resulted in a rise in cytosolic calcium levels, as indicated by Videos 1 & 2 and Figure 4 D (bottom, red trace). Notably, the observed increase in [Ca^2+^]_i_ was CatSper-independent as assured by preincubation (≥ 1 min) with and co-application of the irreversible CatSper inhibiter NNC 55-0396 (Fig. 4D, bottom). Application of NNC 55-0396 alone did serve as a negative control and had no effect on cytosolic calcium levels (Fig. 4 D, black trace). Furthermore, no effects were observed by 1 μM and 10 μM capsaicin (EC_50_ = 711.9 nM [27]) or 500 μM carvacrol (EC_50_ = 490 μM [28]) (Fig. 4 - Suppl. Fig. 1). We therefore conclude that human spermatozoa do not express functional TRPV1 or TRPV3 channels. Instead, our results indicate that the temperature-activated cation channel TRPV4 is functionally expressed and likely provides membrane depolarization in human sperm.

**Figure 4:**
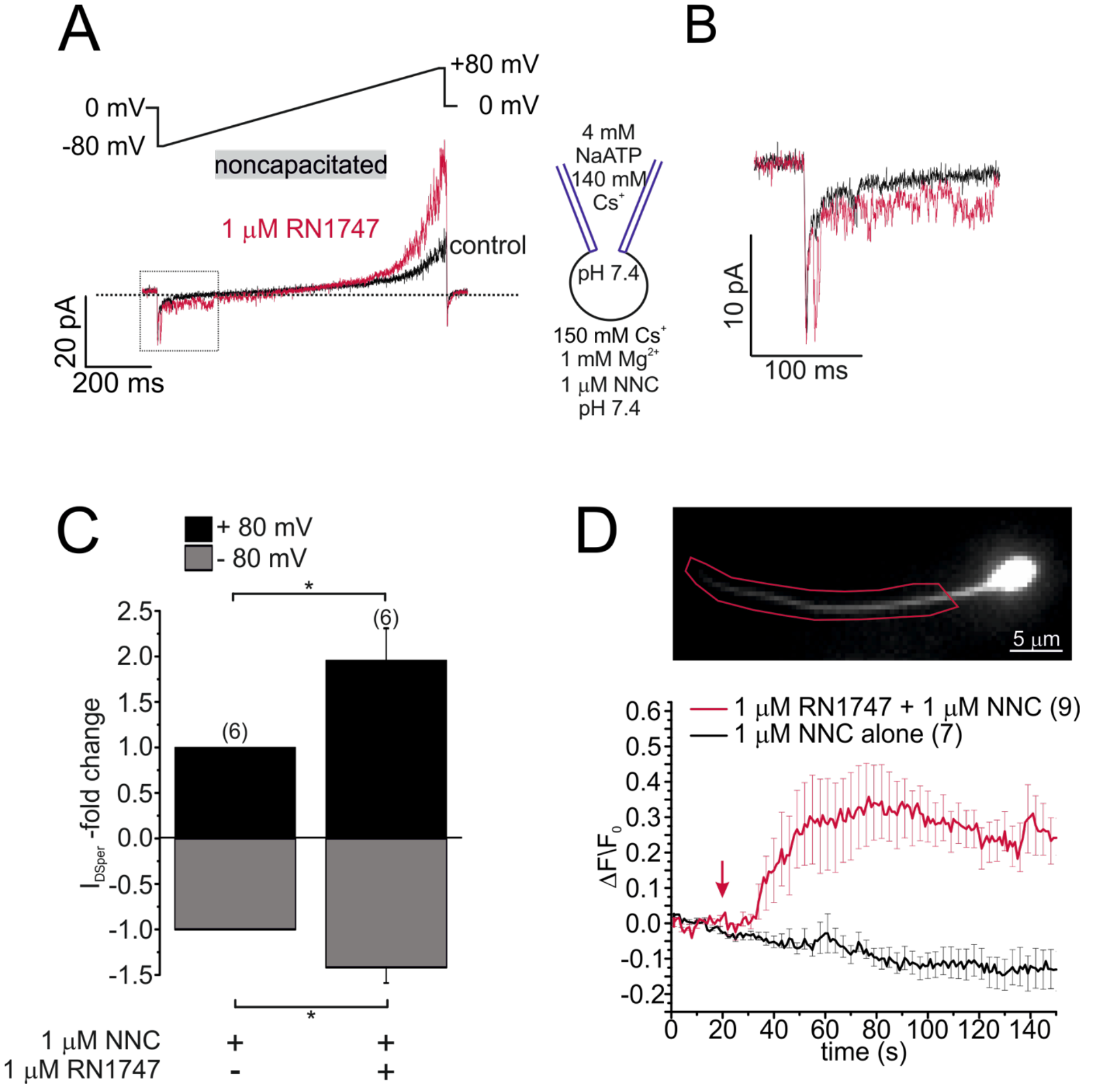
DSper is activated by the TRPV4 agonist RN1747. Original current traces from representative whole-cell patch-clamp recordings of noncapacitated human spermatozoa. Inward- and outward currents were elicited with voltage ramps as depicted. DSper monovalent caesium currents (black trace) are increased after stimulation with 1 μM RN1747 (red trace). Both conditions in presence of 1 μM NNC. (B) Inset of (A) emphasizes how the agonist RN1747 increases DSper’s open probability. (C) Quantification of normalized DSper currents under control conditions and after stimulation with RN1747. Both inward and outward currents show a significant gain upon stimulation with the TRPV4 agonist (factor 1.42 ± 0.17, p = 0.0298 for inward currents, factor 1.95 ± 0.35, p = 0.0209 for outward currents, unpaired t-test, n = 6). Statistical significance (unpaired t-test) was indicated by: \**p ≤0.05*. No variation between human donors were noticed. (D) Noncapacitated spermatozoa were bulk loaded with the calcium indicator fluo-4/AM (top) and fluorescence changes restricted to the flagellar principal piece were analysed upon stimulation with RN1747 (red trace, bottom). Time point of agonist application is indicated by the arrow. In presence of the CatSper inhibitor NNC 55-0369 (and additional preincubation of at least 1 min before agonist application), RN1747 induced a noticeable rise in cytosolic calcium levels. This effect was absent when the cells were stimulated with NNC 55-0369 alone (black trace).

Supporting our functional data, TRPV4 was additionally detected in human sperm on both mRNA and protein levels. Reverse transcriptase PCR performed with a full-length TRPV4 primer pair and mRNA isolated from swim-up purified spermatozoa (Fig. 4 - Suppl. Fig. 2 A) detected a band at the expected size but was absent in negative controls (no reverse transcriptase and no template). Sequencing the isolated PCR product of that specific band (dotted square), yielded the full-length sequence of TRPV4 isoform A (2620 bp, 98 kDa, Q9ERZ8). Moreover, the presence of TRPV4 protein was confirmed by western blotting (Fig. 4 - Suppl. Fig. 2 B). Immunoreactive bands were detected at ~115 kDa in extracts from human testicular tissue (1), capacitated (2) and noncapacitated (3) spermatozoa (Fig. 4 - Suppl. Fig. 2 B). Importantly, when TRPV4 was cloned from human sperm mRNA extracts and recombinantly expressed in HEK293 cells, a band of similar molecular weight was detected (Fig. 4 - Suppl. Fig. 2 C). Finally, immunostaining with anti-hTRPV4 specific antibodies (Fig. 4 - Suppl. Fig. 2 D) yielded an immunopositive signal in the acrosome and flagellum.

## Discussion

Sperm transition to hyperactivated motility is essential for fertilization. Hyperactivation provides the propulsion force required to penetrate through viscous luminal fluids of the female reproductive tract and protective vestments of the egg. The CatSper channel is a key player in the transition to hyperactivated motility [29]. However, proper CatSper function requires three concurrent activation mechanisms: 1) membrane depolarization [6], 2) intracellular alkalization via Hv1-mediated proton extrusion [11], and 3) abundance of progesterone [6], [7]. While the two latter mechanisms have been described in detail, the source of membrane depolarization remained puzzling.

In human spermatozoa, K^+^, Ca^2+^, Cl^−^, and H^+^ conductances have been described [6], [11], [13], [14], [30], [31]. However, the Na^+^ conductance of sperm remained unknown. Upon ejaculation, mammalian spermatozoa are exposed to increased [Na^+^] (~30 mM in cauda epididymis *versus* 100–150 mM in seminal plasma). In the female reproductive tract, Na^+^ levels are similar to those in serum (140–150 mM) [22], [32]. Hence, Na^+^ is ideally suited to provide a depolarizing charge upon sperm deposit into the female reproductive tract.

Here, we record a novel CatSper-independent cation conductance that exhibits outward rectification as well as potentiation upon capacitation. We propose that this novel conductance is carried by the hypothetical “Depolarizing Channel of Sperm” DSper and provides the necessary cation influx for membrane depolarization. *I*_DSper_ is activated by warm temperatures between 22 and 37 °C (Fig. 2, 3 D-E) which makes the protein thermoresponsive to physiologically relevant temperatures (34.4°C in the epididymis [33], 37 °C body core temperature at the site of fertilization). Previous studies showed that capacitated rabbit and human sperm cells have an inherent temperature sensing ability [34], which could be an additional driving force to guide male gametes from the reservoir towards the warmer fertilization site. It is thus very likely, that human spermatozoa express a temperature-activated ion channel, which operates in the described temperature range and enables thermotaxis. The temperature response profile of DSper conforms with previously reported temperature sensitivity of TRPV4 [35], [36]. Moreover, we observed *I*_DSper_ potentiation as well as flagellar Ca^2+^ elevations by the selective TRPV4 agonist RN1747 (Fig. 4).

According to our model (Fig. 5), human spermatozoa are exposed to an increase in both temperature and [Na^+^] upon ejaculation. TRPV4-mediated Na^+^ influx induces membrane depolarization, which in turn activates both Hv1 and CatSper. H^+^ efflux through Hv1 promotes intracellular alkalization and thus enhanced CatSper activation. Approaching the egg, sperm is exposed to P4 and the endocannabinoid anandamide (AEA), both secreted by cumulus cells. P4 binding to ABHD2 releases CatSper inhibition [9] while AEA was shown to activate Hv1 [11]. The resulting opening of CatSper generates a Ca^2+^ wave that propagates along the flagellum and serves as the trigger for hyperactivation. P4 not only potentiates CatSper, it also inhibits KSper-mediated hyperactivation, which gives the CatSper activation cascade an additional impulse [13].

**Figure 5:**
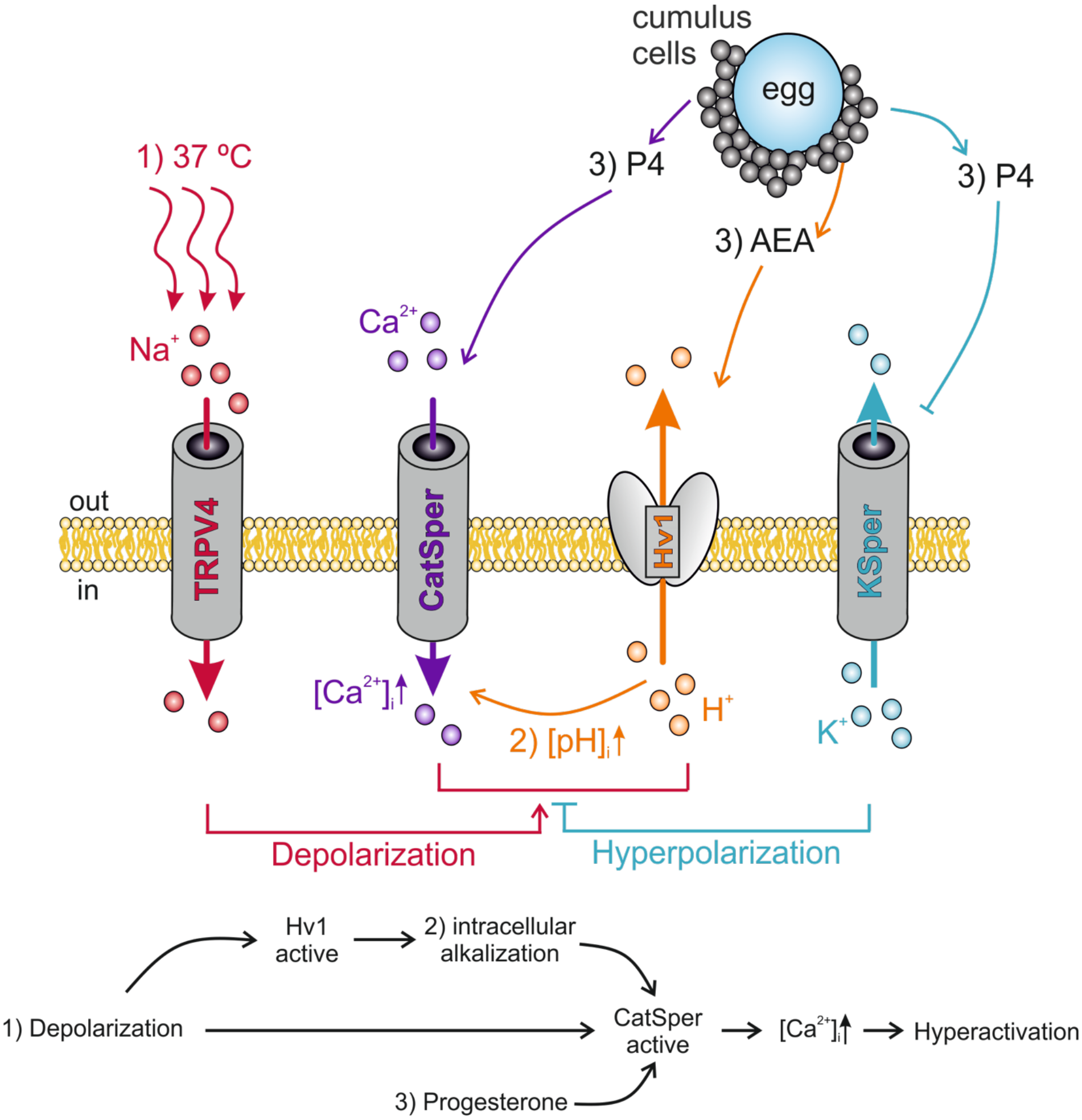
Interdependency of ion channel complexes in the sperm flagellum. Transition into hyperactivated motility is triggered by a CatSper-mediated rise in cytosolic calcium levels. Proper CatSper function requires three concurrent activation mechanisms: 1) membrane depolarization, 2) intracellular alkalization via Hv1-mediated proton extrusion, and 3) abundance of progesterone. In our proposed model the sperm’s sodium channel TRPV4 is activated by warm temperatures (37 °C at the site of fertilization). TRPV4-mediated sodium influx induces 1) membrane depolarization, which in turn activates both Hv1 and CatSper. Hv1 then extrudes protons out of the sperm, thereby leading to 2) intracellular alkalization and further activation of CatSper. Cumulus cells surrounding the egg secrete 3) P4 and AEA. P4 releases CatSper inhibition and blocks KSper-mediated hyperactivation. AEA was shown to potentiate Hv1. The resulting opening of CatSper generates a Ca^2+^ wave that serves as the trigger for hyperactivation.

Using a CatSper2-deficient infertile patient, no remaining cation current was recordable, when both Hv1 and KSper were blocked [37]. However, these recordings were performed in a condition where ATP was absent from the pipette solution. According to Phelps *et al.* intracellular ATP binding to the N-terminal ankyrin repeat domain of TRPV4 has a profound sensitizing effect [38]. Indeed, addition of 4 mM ATP to the pipette solution, allowed us to consistently record TRPV4 activity from fertile human sperm.

Our data suggests that TRPV4 activity is strongly increased upon capacitation. Since capacitation encompasses changes in the phosphorylation state of many proteins [39], and TRPV4 requires tyrosine phosphorylation to function properly [40], it is likely that TRPV4 phosphorylation is required. It would also explain, why only capacitated human spermatozoa appear to be thermotactically responsive [34]. Interestingly, we also observed different *I*_DSper_ kinetics (i.e., less outward rectification) after capacitation. This finding could also be the result of phosphorylation, modified lipid composition or even formation of TRPV4/X heteromers upon capacitation. These aspects will be addressed in future studies.

Selective anti-hTRPV4 antibodies located TRPV4 at the flagellum and acrosome of human sperm (Fig. 4 – Suppl. Fig. 2 D). The localization of TRPV4 in the acrosome region should be evaluated critically, since this compartment is highly antigenic and attracts antibodies in general [41]. However, TRPV4 appears to be distributed in the sperm flagellum. The principal piece of the sperm tail is also the compartment where CatSper and Hv1 reside [6], [11], bringing those three interdependent ion channels in close proximity to each other.

TRPV4 – more precisely its hyperfunction - might underlie the aversive effect of increased scrotal temperatures on sperm production and epididymal preservation. As proposed by Bedford *et al.*, increased scrotal temperatures when clothed contribute substantially to the inferior quality of human ejaculate [42]. By contrast, TRPV4 might represent an attractive target for male fertility control, since TRPV4 is likely to lie upstream in the signaling cascades leading to sperm hyperactivation and can be heterologously expressed for high-throughput functional studies.

## Materials and methods

### Human sperm cells

A total of 3 healthy male volunteers were recruited to this study, which was conducted with approval of the Committee on Human Research at the University of California, Berkeley (protocol 10-01747, IRB reliance #151). Informed consent was obtained from all participants. Ejaculates were obtained by masturbation and spermatozoa were purified following the swim-up protocol as previously described [6]. In-vitro capacitation was accomplished by 4-hour incubation in 20 % Fetal bovine serum, 25 mM NaHCO_3_ in HTF buffer [11] at 37 °C and 5 % CO_2_.

### Reagents

NNC 55-0396 and RN 1747 was purchased from Tocris Bioscience (Bristol, UK), capsaicin from Cayman Chemical (Ann Arbor, USA), fluo-4/AM is from Invitrogen (Thermo Fisher Scientific, Carlsbad, USA) and all other compounds were obtained from Sigma (St. Louis, USA).

### Electrophysiology

For electrophysiological recordings, only the ultra-pure upper 1 ml of the swim-up fraction was used. Single cells were visualized with an inverse microscope (Olympus IX71) equipped with a differential interference contrast, a 60 × Objective (Olympus UPlanSApo, water immersion, 1.2 NA, **∞**/0.13-0.21/FN26.5) and a 1.6 magnification changer. An AXOPATCH 200B amplifier and an Axon^TM^ Digidata 1550A digitizer (both Molecuar Devices, Sunnydale, CA, USA) with integrated Humbug noise eliminator was used for data acquisition. Hardware was controlled with the Clampex 10.5 software (Molecular Devices). We monitored and compensated offset voltages and pipette capacitance (C_fast_). Gigaohm seals were established at the cytoplasmic droplet of highly motile cells in standard high saline buffer (“HS” in mM: 135 NaCl, 20 HEPES, 10 lactic acid, 5 glucose, 5 KCl, 2 CaCl_2_, 1 MgSO_4_, 1 sodium pyruvate, pH 7.4 adjusted with NaOH, 320 mOsm/l) [43], [17]. The patch pipette was filled with 140 mM CsMeSO_3_, 20 mM HEPES, 10 mM BAPTA, 4 mM NaATP, 1 mM CsCl (pH 7.4 adjusted with CsOH, 330 mOsm/l). For recordings from capacitated spermatozoa, BAPTA was substituted for 5 mM EGTA and 1 mM EDTA. Transition into whole-cell mode was achieved by applying voltage pulses (499–700 mV, 1-5 ms, V_hold_ = 0 mV) and simultaneous suction. After establishment of the whole-cell configuration, inward and outward currents were elicited via 0.2 Hz stimulation with voltage ramps (−80 mV to +80 mV in 850 ms, V_hold_ = 0 mV, total 1000 ms/ramp). Data was not corrected for liquid junction potential changes. To ensure stable recording conditions, only cells with baseline currents (in HS solution) ≤ 10 pA at −80 mV were used for experiments. Under “HS” condition, CatSper and DSper currents were considered to be minimal, thus any remaining baseline current represented the cells leack current. During whole-cell voltage-clamp experiments, the cells were continuously superfused with varying bath solutions utilizing a gravity-driven perfusion system. If not stated otherwise, electrophysiological experiments were performed at 22°C. Temperature of the bath solution was controlled and monitored with an automatic temperature control (TC-324B, Warner Instrument Corporation, Hamden, CT, USA). Both, CatSper and Dsper currents were recorded under symmetric conditions for the major permeant ion. Under these conditions, the bath solution was divalent free (“DVF”) containing (in mM) 140 CsMeSO_3_, 20 HEPES, 1 EDTA, and pH 7.4 was adjusted with CsOH, 320 mOsm/l. To isolate Dsper conductances, monovalent currents through CatSper channels were inhibited by supplementing the DVF solution with 1 mM Mg^2+^ [6]. Experiments with different bath solutions were performed on the same sperm cell. Signals were sampled at 10 kHz and low-pass filtered at 1 kHz (Bessel filter; 80 dB/decade). Pipette resistance ranged from 9 – 15 MΩ, access resistance was 21–100 MΩ, membrane resistance ≥ 1.5 GΩ. Membrane capacitance was 0.8-1.3 pF and served as a proxy for the cell surface area and thus for normalization of current amplitudes (i.e., current density). Capacitance artifacts were graphically removed. Statistical analysis was done with Clampfit 10.3 (Molecular Devices, Sunnyvale, CA, USA), OriginPro 8.6 (OriginLab Corp., Northampton, MA, USA) and Microsoft Excel 2016 (Redmond, WA, USA). Statistical data are presented as mean **±** standard error of the mean (SEM), and (n) indicates the number of recorded cells. Statistical significance was determined with unpaired t-tests.

### Calcium Imaging

All calcium imaging experiments were performed in HS solution. Prior to fluorescence recording, swim-up purified human spermatozoa [29] were bulk loaded with 9 μM fluo-4/AM (dissolved in DMSO) and 0.05% Pluronic (dissolved in DMSO) in HS solution for 30 min at room temperature. Cells were then washed with dye-free HS solution and allowed to adhere to glass imaging chambers (World Precision Instruments, Sarasota, USA) for 1 min. Via continuous bath perfusion, the attached spermatozoa were presented with alternating extracellular conditions (HS +/− agonist/antagonist). Fluorescence was recorded at 1 Hz, 100 ms exposure time over a total time frame as indicated. Imaging was performed using a Spectra X light engine (Lumencore, Beaverton, USA) and a Hamamatsu ORCA-ER CCD camera. Fluorescence change over time was determined as ΔF/F_0_ where ΔF is the change in fluorescence intensity (F - F_0_) and F_0_ is the baseline intensity as calculated by averaging the fluorescence signal of the first 20 s in HS solution. Regions of interest (ROI) were restricted to the flagellar principal piece of each cell by manual selection in ImageJ (Java, Redwood Shores, CA, USA). It should be noted that only a fraction of cells responded to RN1747 stimulation, reflecting the heterogeneity of the human sperm pool. Only responsive cells were considered for statistical analysis. However, 0 % of the imaged cells, showed a response to carvacrol or capsaicin. Statistical data are presented as mean **±** standard error of the mean (SEM), and (n) indicates the number of recorded cells.

### Immunocytochemistry

Purified spermatozoa were plated onto 20-mm coverslips in HS and allowed to attach for 20 min. The cells were fixed with 4% paraformaldehyde (PFA) in PBS for 20 min and washed twice with PBS. Additional fixation was performed with 100% ice-cold methanol for 1 min with two washing steps in PBS. Cells were blocked and permeabilized by 1-hour incubation in PBS supplemented with 5 % immunoglobulin-γ (IgG)–free BSA and 0.1 % Triton X-100. Immunostaining was performed in the same blocking solution. Cells were incubated with primary antibodies (rabbit polyclonal αTRPV4, 1:100, abcam ab39260) overnight at 4°C. After extensive washing in PBS, secondary antibodies (mouse monoclonal αRabbit-DyLight™488, 1:1000, Jackson 211-482-171) were added for 45 min at room temperature. After vigorous washing, cells were mounted with ProLong Gold Antifade with DAPI reagent (Life Technologies, Carlsbad, CA) and imaged with a confocal microscrope.

### RT-PCR and cloning

Total donor-specific RNA was extracted from purified spermatozoa with a QIAGEN RNAeasy mini kit followed by complementary DNA synthesis with a Phusion RT-PCR kit (Finnzymes, MA, USA). The donor-specific translated region of TRPV4 (cDNA) was amplified with the primers forward 5- ACAGATATCACCATGGCGGATTCCAGCG -3’ and reverse 5’-AACACAGCGGCCGCCTAGAGCGGGGCGTCATC-3’, and was subcloned into a pTracer-CMV2 vector (Invitrogen) using the restriction sites: EcoR V and Not I. TRPV4 identity was sequence verified.

### Immunoblotting

The highly motile sperm fraction was separated from other somatic cells (mainly white blood cells, immature germ cells, and epithelial cells) by density gradient consisting of 90% and 50% isotonic Isolate (Irvine Scientific, CA) solution diluted in HS solution with the addition of protease inhibitors (Roche). Protease inhibitors were used throughout the whole procedure. After centrifugation at 300 g for 30 min at 24C, the sperm pellet at the bottom of the 90% layer was collected, diluted ten times, and washed in HS by centrifugation at 2000 g for 20 min. Cells were examined by phase-contrast microscopy for motility and counted before centrifugation. Contamination of the pure sperm fraction by other cell types was minimal, with less than 0.2% of somatic cells, which was below the protein detection threshold for immunoblotting applications. The pellet was subjected to osmotic shock by a 5 min incubation in 0.5x HS solution, the addition of 10 mM EDTA and 10mM dithiothreitol (DTT) for 10 min, and sonication in a water bath at 25 °C for 5 min. Osmolarity was adjusted by addition of 10x phosphate-buffered saline (PBS). Laemmli sample buffer (5x) was added to a final 1x concentration, and the DTT concentration was adjusted to 20 mM. An additional 5 min sonication and boiling at 100 °C for 5 min were performed. The total crude cell lysate was loaded onto a 4%– 20% gradient Tris-HCl Criterion SDS-PAGE (BioRad) with 500,000 sperm cells/well. TRPV4- and empty vector-transfected HEK293 cells were lysed in 2x Laemmli sample buffer and subjected to SDS-PAGE. Ten thousand cells per well were loaded onto SDS-PAGE. After transfer to polyvinylidene fluoride membranes, blots were blocked in 0.1% PBS-Tween20 with 3% IgG-free BSA for 15 min and incubated with primary antibodies overnight at 4 °C. Blots were probed with rabbit anti-b-tubulin antibodies (Abcam), mouse monoclonal anti-actin C4 antibodies (Abcam), or anti-TRPV4 antibodies (Abcam). After subsequent washing and incubation with secondary horseradish peroxidase-conjugated antibodies (Abcam), membranes were developed with an ECL SuperSignal West Pico kit (Pierce).

## Acknowledgements

We thank Dr. Junji Suzuki (UCSF, CA) for the help with calcium imaging. We also thank Dr. Julio F. Cordero-Morales (University of Tennessee Health Science Center, Memphis, TN) for the initial input and suggestions. M.S. is a Lichtenberg Professor of the Volkswagen Foundation and acknowledges support from the FENS-Kavli Network of Excellence. This work was supported by a DAAD fellowship to N.M., and by NIH R01GM111802, R21HD081403, Pew Biomedical Scholars Award, Alfred P. Sloan Award, and Packer Wentz Endowment Will to P.V.L.

---

**Fig. 1 - Supplementary Figure 1:**
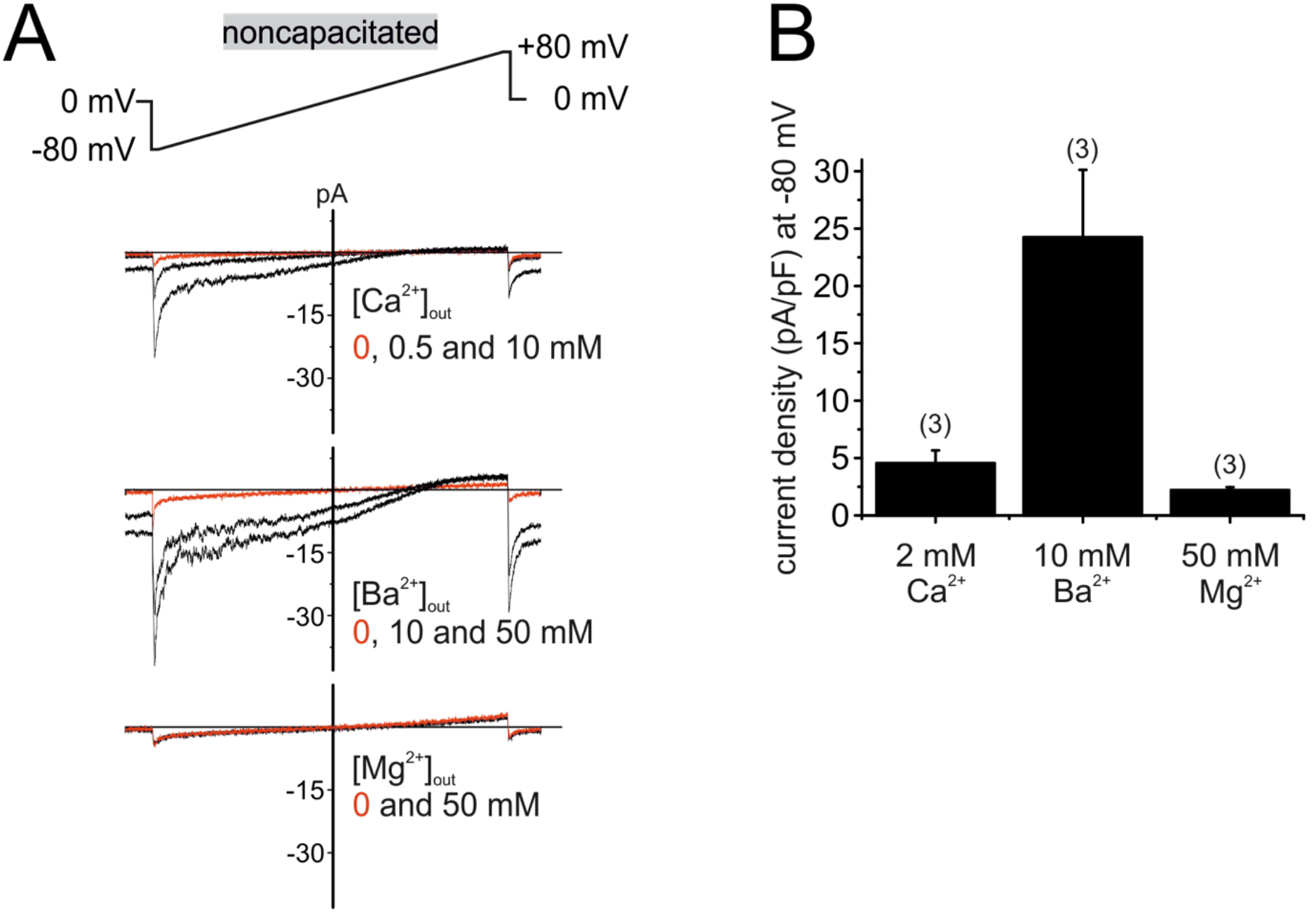
Human CatSper conducts Ca^2+^ and Ba^2+^, but not Mg^2+^. (A) Original current traces from whole-cell voltage-clamp recordings of noncapacitated human spermatozoa. Inward- and outward currents were elicited with voltage ramps as depicted. Pipette solution was: 140 mM NMDG, 100 mM Hepes, 5 mM EGTA, 5 mM EDTA, 330 mosmol, pH 7.3, composition of bath solution was: 500 nM progesterone, 100 mM Hepes, 130 mM NMDG, plus X mM Ca2+, Ba2+ or Mg2+ as depicted, 317 mosmol, pH 7.4. When the major permeable extracellular cation was Ca^2+^ or Ba^2+^, negative membrane potentials induced concentration-dependent inward currents. In the presence of Mg^2+^, CatSper currents remained at baseline level (0 mM), indicating that human CatSper is not permeable for Mg^2+^. (B) Quantification of current densities (pA/pF) for either Ca^2+^, Ba^2+^ or Mg^2+^ inward currents through CatSper.

**Fig. 1 - Supplementary Figure 2:**
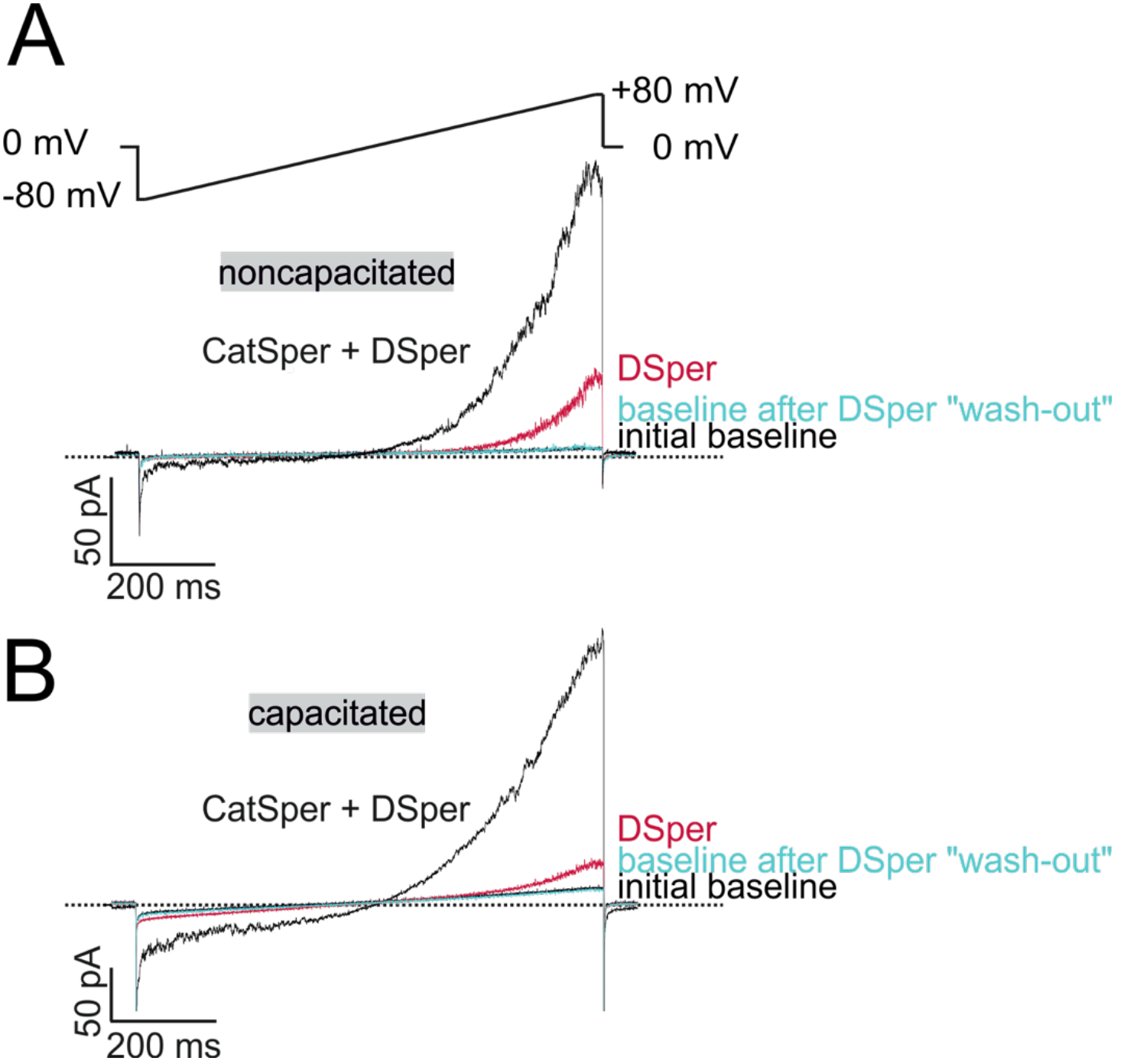
DSper currents were recorded under stable conditions. (A-B) Original current traces from representative whole-cell patch-clamp recordings of noncapacitated (A) and capacitated (B) human spermatozoa. Inward- and outward currents were elicited with voltage ramps as depicted in (A). Represented are three conditions – baseline (in HS solution), CatSper + DSper currents and isolated DSper currents. Whole-cell currents returned to their initial baseline level after returning to HS solution, indicating that the recorded DSper currents are not a remnant of an increased leak-current.

**Fig. 4 - Supplementary Figure 1:**
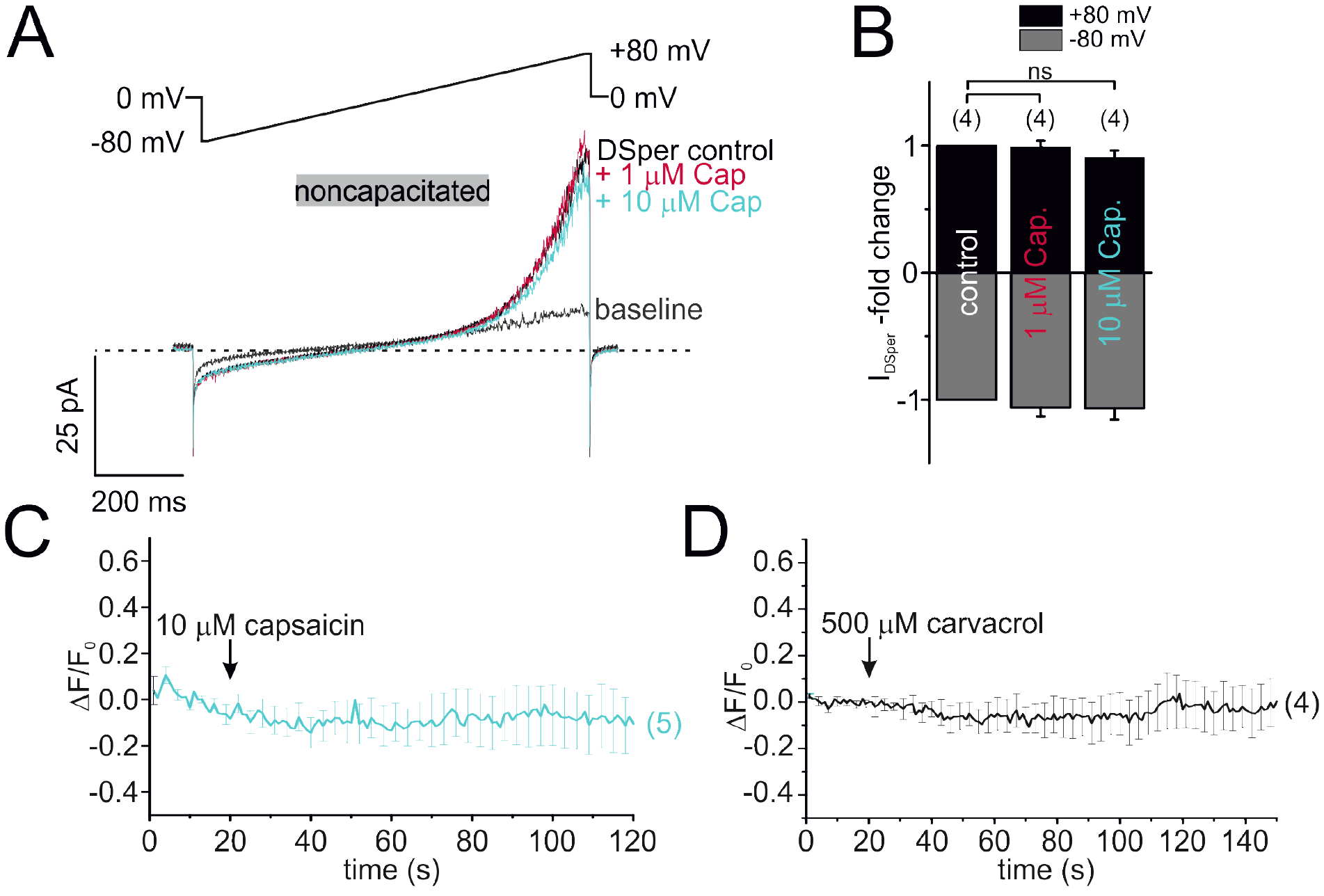
TRPV1 and TRPV3 is not functionally expressed in human spermatozoa. (A) Original current traces from representative whole-cell patch-clamp recordings of noncapacitated human spermatozoa. Inward- and outward currents were elicited with voltage ramps as depicted. Stimulation with two different concentrations (1 and 10 μM) of the specific TRPV1 agonist capsaicin did not induce any significant effect on DSper control inward or outward currents. Quantification of normalized DSper currents w/ and w/o agonist in (B). (C) Single-cell calcium imaging results confirmed our electrophysiological findings. Application of 10 μM capsaicin had no effect on cytosolic calcium levels. (D) Our Single-cell calcium imaging approach did not reveal any notable effect of the TRPV3 specific agonist carvacrol (500 μM).

**Fig. 4 - Supplementary Figure 2:**
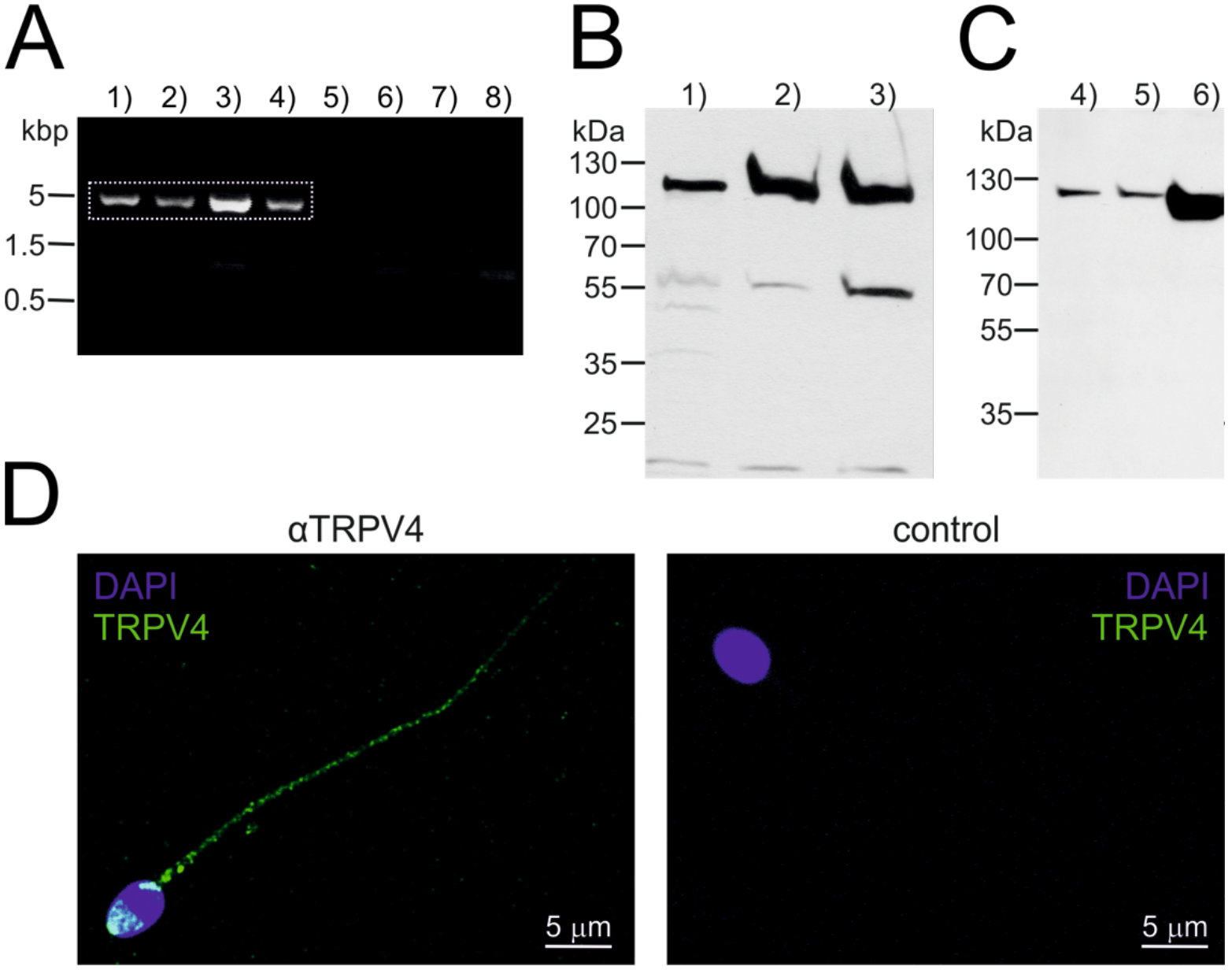
TRPV4 can be detected on protein and mRNA level. (A) RT-PCR using a full-length TRPV4 primer pair, mRNA isolated from swim-up purified noncapacitated spermatozoa and PCR conditions as follows: 1) – 4) varying annealing temperatures (52-60 °C), 5) - 6) negative control in absence of the reverse transcriptase enzyme (Ta = 50 °C, 56 °C), 7) – 8) no template control (Ta = 56 °C). Dotted square marks bands that were selected for gene product sequencing. (B) Western blotting confirms the presence of the TRPV4 peptide in 1) human testicular tissue 2) capacitated and 3) noncapacitated spermatozoa. Immunopositive bands can be detected in all three samples at approx. 115 kDa. (C) TRPV4 was cloned from human sperm and recombinantly expressed in HEK293 cells. Western blotting results are shown for 4) nontransfected HEK293 cells, 5) cells transfected with the empty vector and 6) HEK293 cells transfected with the TRPV4-containing vector. An intense immunopositive band can be detected in line 6), at same hight as in (B 1-3). Weak bands in 4) and 5) suggest endogenous expression of TRPV4 in HEK293 cells. (D) Confocal fluorescence images of immunostainings against TRPV4. (Left) noncapacitated spermatozoa were labeled with an anti-TRPV4 selective antibody and a Dylight488-conjugated sencondary antibody. Nuclear dye DAPI locates the sperm head. Immunopositive fluorescent signal was detected in the sperm flagellum and the acrosome region. (Right) Negative control shows no unselective signal in the absence of the primary antibody, but in presence of the secondary.

